# Recent land use and land cover pressures on Iberian peatlands

**DOI:** 10.1101/2023.07.03.547480

**Authors:** Raquel Fernandes, Miguel Geraldes, Elizabete Marchante, Jorge Durán, César Capinha

## Abstract

Iberian peatlands have been severely affected by land use and land cover (LULC) changes. Despite these pressures, some peatlands persist in the region, although their susceptibility to LULC change remains poorly understood. This study presents the most detailed and extensive distribution data for Iberian peatlands to date and analyzes the dynamics and drivers of LULC in Iberian peatlands and their surrounding areas. We compiled peatland records from various sources and used Corine Land Cover Change layers to determine LULC shifts for 1990, 2000, 2006, 2012, and 2018. Environmental and socioeconomic variables were used to create Boosted Regression Tree models explaining spatial variations in the mean percentage of changed area. Analysis of 270 peatland locations in the Iberian Peninsula revealed that forests and seminatural areas constituted over 80% of the peatland’s surroundings. Agricultural areas expanded the most, except between 2006 and 2012 when the artificial areas showed more gains. While most areas experienced an average change of 0%-9.51% of the total area, between 1990 and 2018, lowland peatlands (littoral and sublittoral) suffered more intense changes (9.51% to 38.43%). Our models showed that only elevation and agricultural area density were relevant predictors of spatial distribution changes. Upland Iberian peatlands showed lower susceptibility to LULC changes, while lowland peatlands underwent remarkable transformations. This study substantially expands previous knowledge about the distribution and conservation needs of these ecosystems in the Iberian Peninsula, especially those in littoral and sublittoral lowlands.

## 1. Introduction

Peatlands store the largest amounts of carbon (C) per unit area (Temmink et al. 2022), an equivalent of about 25% of the world’s soil C (Parish et al. 2008). When natural conditions are disturbed (e.g., abnormally low water table level or vegetation removal), peat dries and oxides due to the contact with atmospheric oxygen, leading to large C losses to the atmosphere, mainly in the form of carbon dioxide (CO_2_) (Parish et al. 2008, Joosten et al. 2012).

Land use and land cover (LULC) changes are one of the major degradation drivers of these carbon-rich ecosystems, particularly intentional drainage (Tanneberger, Appulo, et al. 2021), resulting in total CO_2_ emissions from disturbed peatlands accounting for approximately a quarter of the total emissions associated with the land use, land-use change, and forestry sector globally (Tubiello et al. 2016). For example, boreal peatlands have been extensively modified for peat extraction and agriculture, which dates back at least to the 19th century in several European countries (Jansen et al. 2009; Holden 2017; Tanneberger et al. 2020; Sinyutkina 2021). Similarly, temperate peatlands have been drained for agricultural land expansion or to reduce the risk of malaria transmission (Evrendilek et al. 2011; Joosten et al. 2017; Sánchez-Espinosa & Schröder 2019). Peatlands have also undergone rapid degradation in tropical regions, driven by agricultural expansion and fires (Miettinen et al. 2016; Hergoualc’h et al. 2017; Sutikno et al. 2020).

Peatlands in the Iberian Peninsula have been described as rare, or with low peat accumulation rates owing, to a large extent, to dry and warm climate conditions in most of the Peninsula (Tanneberger Moen et al. 2021; Pontevedra-Pombal et al. 2017). Furthermore, peatlands in this region have also been experiencing strong pressure in recent decades from multiple human activities, such as drainage for both farmland conversion and malaria-risk reduction, overgrazing, afforestation, peat extraction for horticulture and the construction of wind farms (Heras Pérez et al. 2017; Mateus et al. 2017; Chico et al. 2019). In the most severe cases, these ecosystems have completely disappeared (Tanneberger, Moen, et al. 2021). Some of these pressures could be partially attributed to the adoption of the EU Common Agricultural Policy (CAP) by Portugal and Spain, the two countries comprising most of the Peninsula, after they became members of the European Union (EU) in 1986 (Jones et al, 2011). Rural abandonment was also widespread in many regions of the Peninsula following EU membership (Jones et al. 2011). While this led to a decline in agricultural activities, potentially reducing this source of pressure on peatlands, it has also led to uncontrolled plant succession (natural or by invasive plants), and alteration and loss of habitats, ultimately affecting the maintenance of the particular conditions required by these ecosystems (Plieninger 2006; Lasanta & Vicente-Serrano 2012). Yet, despite the pressures, several Iberian peatland areas persisted, mainly in northern mountain areas with high precipitation, but also in low-energy wetlands (e.g., river floodplains and estuaries and freshwater coastal lagoons) (Joosten et al. 2017; Pontevedra-Pombal et al. 2017). However, recent LULC dynamics and consequent impacts on the degradation of these ecosystems are not well known in this region. These peatlands provide important ecosystem services, such as carbon storage and climate regulation services (Martínez-Cortizas et al. 2000), or habitat for rare and priority species provision (Mateus et al. 2017; Neto et al. 2021). Furthermore, they are under various types of protection status (e.g., Protected Areas and Classified Areas) (Mateus et al. 2017) and are considered habitats of Community interest under the Habitats Directive 92/43/EEC (European Commission 2023) and central to achieving the carbon emission reduction targets of the Paris Agreement (Minasny et al. 2019).

Therefore, understanding LULC dynamics in the remaining Iberian peatlands is crucial to support management and conservation measures. In this study, we addressed these knowledge gaps by providing the most comprehensive distribution data on Iberian peatlands to date. We also analyzed the recent LULC dynamics in their surrounding areas to understand how LULC has impacted these ecosystems in recent decades. Further, to provide critical insights into peatlands’ current conservation needs across the Iberian Peninsula, our analysis included the measurement of temporal trends of change, LULC transition dynamics, and the testing for spatially explicit drivers of the identified patterns.

## 2. Methods

### 2.1. Study area

The Iberian Peninsula, the study area (Fig.1), is the westernmost region of Europe, located between 36°00’N and 43°47’N and 9°29’ W and 3°19’ E and is surrounded by the Mediterranean Sea, the Cantabrian Sea, and the Atlantic Ocean. Around two-thirds of the total surface area is covered by plains (the Mesetas), surrounded by mountain chains, with the whole Peninsula reaching a mean altitude of 600 m a.s.l. (Pontevedra-Pombal et al. 2017). It is located in a climatic transition belt. While the Mediterranean climate is dominant in its core and in the south, the temperate oceanic climate occupies the northern fringe (Kottek et al. 2006; Heras Pérez et al. 2017; Mateus et al. 2017). In addition, some continentality and mountain effects can influence local climates.

**Fig. 1.**
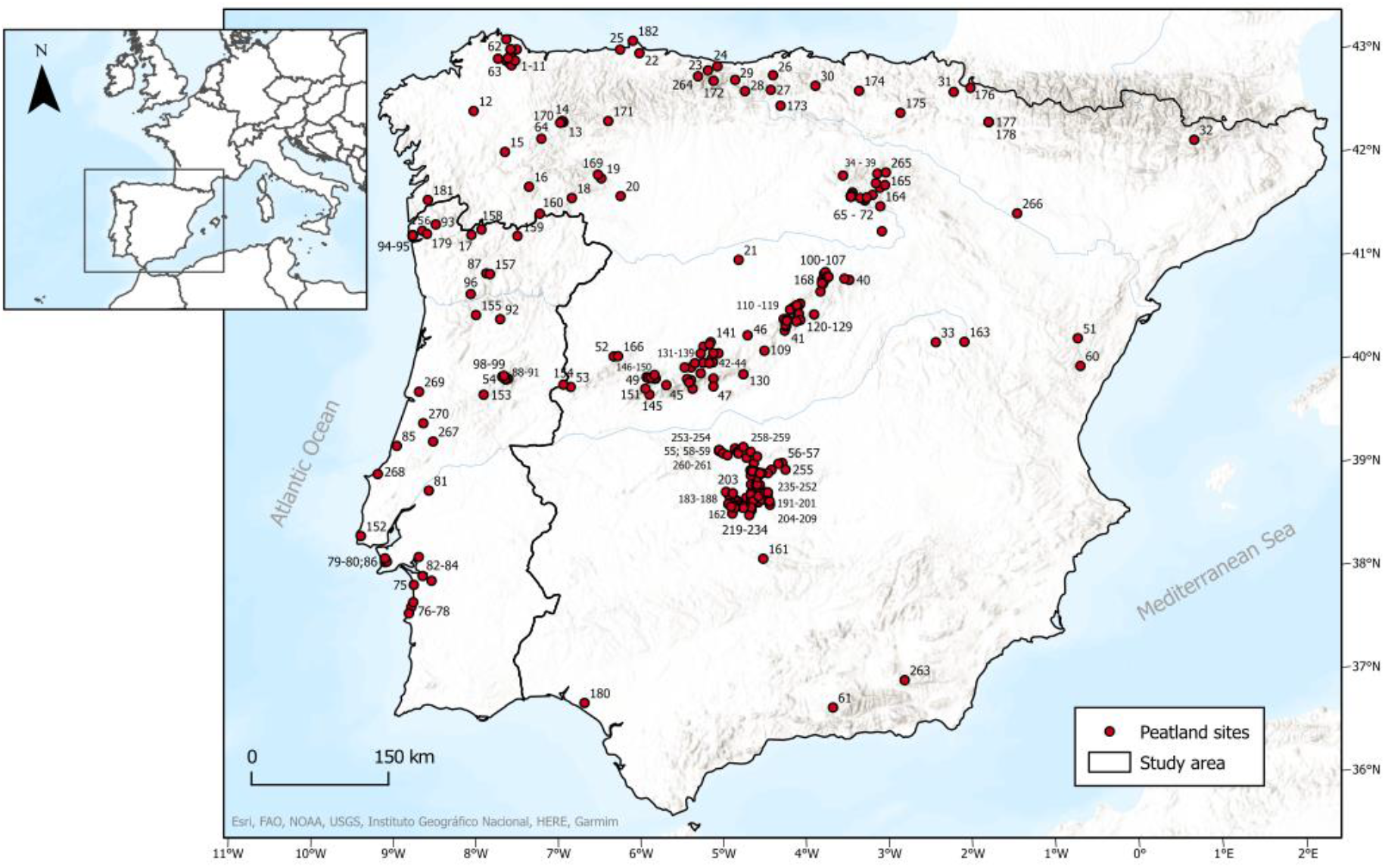
Distribution of the 270 peatland sites (red circles) in the Iberian Peninsula, based on literature and fieldwork. The information about each peatland can be found in the Supplemental Information Table 1

**Table 1.** Description of the calculated measures from the cross-tabulation matrices. *T1* represents the initial time (“from”), and *T2* represents the final time (“to”)

| Measure | Description |
| --- | --- |
| Transition in the same class | The proportion of land cover that had a transition within the same class (e.g., “Coniferous forest” convert to “Transitional woodland/shrub”, both within the same level 1 class “Forest and seminatural areas”). |
| Transition to another class | The proportion of land cover that had a transition to other classes. |
| Gain | The proportion of the landscape that experienced gross gain is the difference between the total of $T2$ and the transition in the same class. |
| Loss | The proportion of the landscape that experienced gross loss is the difference between the total of $T1$ and the transition in the same class. |
| Total change | Sum of the gain and loss. |

Regarding the European Mire region distribution, Portugal and Spain share the western rims of three mire regions the Southern European marsh region, which covers most Iberian Peninsula area; the Atlantic bog region, along the oceanic façade of western Europe; and the Central and southern European mountain compound region, in the mountain areas (Joosten et al. 2017; Tanneberger Moen et al. 2021).

### 2.2. Iberian peatlands database

Despite recent efforts to compile distribution data on Iberian peatlands (e.g., Heras Pérez et al. (2017), Mateus et al. (2017), Pontevedra-Pombal et al. (2017)), the criteria used often do not reflect the wider distribution of peatlands in the Iberian Peninsula (e.g., peatlands with specific international importance to science, or peatland definitions used in boreal latitudes). Therefore, in this work, the reference to “peatlands” encompasses peat-related ecosystems like bogs, transition mires, and fens. We assembled an up-to-date geographical database of Iberian peatland sites, starting with a review of online literature, using Google Scholar and Google Search. We used multiple search terms (“turba”, “turbera”, “turfa”, “turfeira”, “peat”, “peatland” and “mire”) and searched published and unpublished scientific articles, doctoral theses, books, and technical reports, from different research areas (e.g., paleoenvironment, palynology, paleoecology). To ensure the inclusion of currently existing peatlands, we focused on the most up-to-date Iberian peatland maps available (Heras Pérez et al. 2017; Mateus et al. 2017; Pontevedra-Pombal 2017) and literature published from 2010 onward. In addition, we also included a set of 35 records of sites identified during fieldwork performed by one of the authors (MG), mostly during 2011-2015 and 2018-2021. We excluded records of buried peat and peat at the surface after erosion (i.e., former active peatland ecosystems), in order to consider peaty topsoils only. Records with coordinates with less than 10 m of spatial precision were also not incorporated, to make sure only accurate locations were included. For each site, we collected the following information: altitude, country, morpho-structural unit/subunit, and type of peatland (lowland or upland, using the upland-lowland divide after Ramos & Ramos-Pereira (2020) and Lindsay (2016). This information was retrieved from the source publications when available. If not available, elevation data were extracted from the Digital Elevation Model derived from the SRTM 1 Arc-Second Global elevation data (EROS, 2018), using GIS software ArcGIS Pro 3.1.1.. Morpho-structural information was complemented with related literature (Pereira et al. 2014; Alarcón & Cunha 2019; Vergés & Kullberg 2019; Ramos & Ramos-Pereira 2020).

### 2.3. Land use land cover change analysis

#### 2.3.1. Data

To assess LULC dynamics in the areas surrounding each peatland, we used the CORINE Land Cover (CLC) products from the Copernicus Land Monitoring Service (Büttner et al. 2021). Portugal and Spain have their own LULC cartography, the Portuguese “Carta de Ocupação dos Solos” (Direção-Geral do Território 2022), and the Spanish SIOSE/High Resolution SIOSE (Instituto Geográfico Nacional 2022). However, these datasets do not allow a homogeneous land cover classification and the temporal matching of available information for both countries, contrary to CORINE products.

First, we performed a general description of the LULC available classes of the studied areas using the CLC status layers, using the available years of 1990, 2000, 2006, 2012, and 2018. The CLC status is 100 m resolution and uses a Minimum Mapping Unit (MMU) of 25 ha for aerial phenomena (Büttner et al. 2021). To perform these assessments, we aggregated the 44 level 3 classes based on CLC level 1, comprising five classes: artificial surfaces, agricultural areas, forest and seminatural areas, wetlands, and water bodies. A 2000 m radius buffer around each site was designed, for which we measured the percentage of area covered by each class. This radius value is considered appropriate since it covers the known width of Spanish peatlands, which can extend up to 400 ha (Heras Pérez et al. 2017), and includes peatlands in Portugal, based on previous mappings that used an 800 m radius (Tanneberger, Moen, et al. 2021). Moreover, the purpose of the buffer areas is also to encompass the surrounding landscape of peatlands to analyze human-mediated dynamics in LULC. Despite the assumed adequacy of this buffer radius, we performed the same set of analyses with radius lengths of 1000 m, and 4000 m, to assess the robustness of results to this parameter. In all cases, the buffers do not encompass oceanic areas.

Second, we used CLC-Change vector data layers (CHA) to assess the LULC class transitions between two contiguous status assessments (1990-2000, 2000-2006, 2006-2012, and 2012-2018). The CHA are recommended to assess LULC changes, rather than directly from CLC status, since CHA comprises LULC changes with 5 ha of MMU (Büttner et al. 2021; García Álvarez & Camacho Olmedo 2023). As for the previous analyses, these measurements were made using the five, level 1, CLC classes. The CHA vector layers were reclassified into raster layers, with the “from” class and “to” class information for the above-mentioned periods. Based on this information, we calculated a cross-tabulation matrix of LULC transitions, adapting the methodology proposed by Pontius et al. (2004). The diagonal values of the matrix represent a change within the same class, from the initial time (*T1*) to the final time (*T2*), and the non-diagonal values are relative to the transitions between different classes. From these measurements, we calculated the percentage of LULC transitions in the same class and to another class, the amount of gross gain and gross loss of each class, and the total change (Table 1).

#### 2.3.2. Temporal trends

Non-parametric Kruskal-Wallis tests (Ostertagová et al. 2014) were performed to test whether the distribution of the percentage of CLC change of each site for each period was the same (null hypothesis, H0) or distributed differently (alternative hypothesis, H1). The Kruskal-Wallis test does not identify which period(s) differ from others if existing, so we performed the post-hoc Dunn tests (Dinno 2015) to identify which period(s) differ from others. The Kruskal-Wallis tests were performed in R, using the “Kruskal.test()” function, and the Dunn tests was calculated with the “FSA” R package (Ogle et al. 2023).

### 2.4. Model of LULC pressures over peatlands

#### 2.4.1. Spatial drivers of CLC dynamics

To unveil the main drivers of spatial differences in LULC dynamics across the Peninsula, we performed a multivariate analysis assessing the influence of specific spatial variables. The response variable represented the mean changed area (%), between 1990 and 2018, which is relative to the LULC dynamic propensity during the study period, hence acting as an indicator of LULC pressure on peatlands. To explain the spatial variation in this indicator, we considered nine spatial variables, representing environmental and socioeconomic factors (Table 2). Environmental variables (Table 2) were considered because peatlands located at higher elevations and slopes as well as in harsh climates are hypothesized to have less LULC variation, for example in high mountain areas, these ecosystems may not be reclaimed for agriculture or other land uses (Mateus et al. 2017). The socioeconomic group (Table 2) aims to represent the impact of human distribution and activities. Overall, we expect higher LULC dynamics (and presumed pressures over peatlands) where road density is higher (Millington et al. 2007; Xie et al. 2021), as well as in areas with higher levels of economic activity (Sica et al. 2016). The percentage of density of agricultural areas was calculated based on the ratio between the area occupied by the “agricultural areas”, from CLC status layers, and the total area of each buffer. In addition, we included the variable “country” distinguishing whether the peatland was located in Portugal or Spain to consider the potential influence of national-level effects, such as distinct land-use or environmental policies and agricultural practices (Naimi 2022).

**Table 2.**
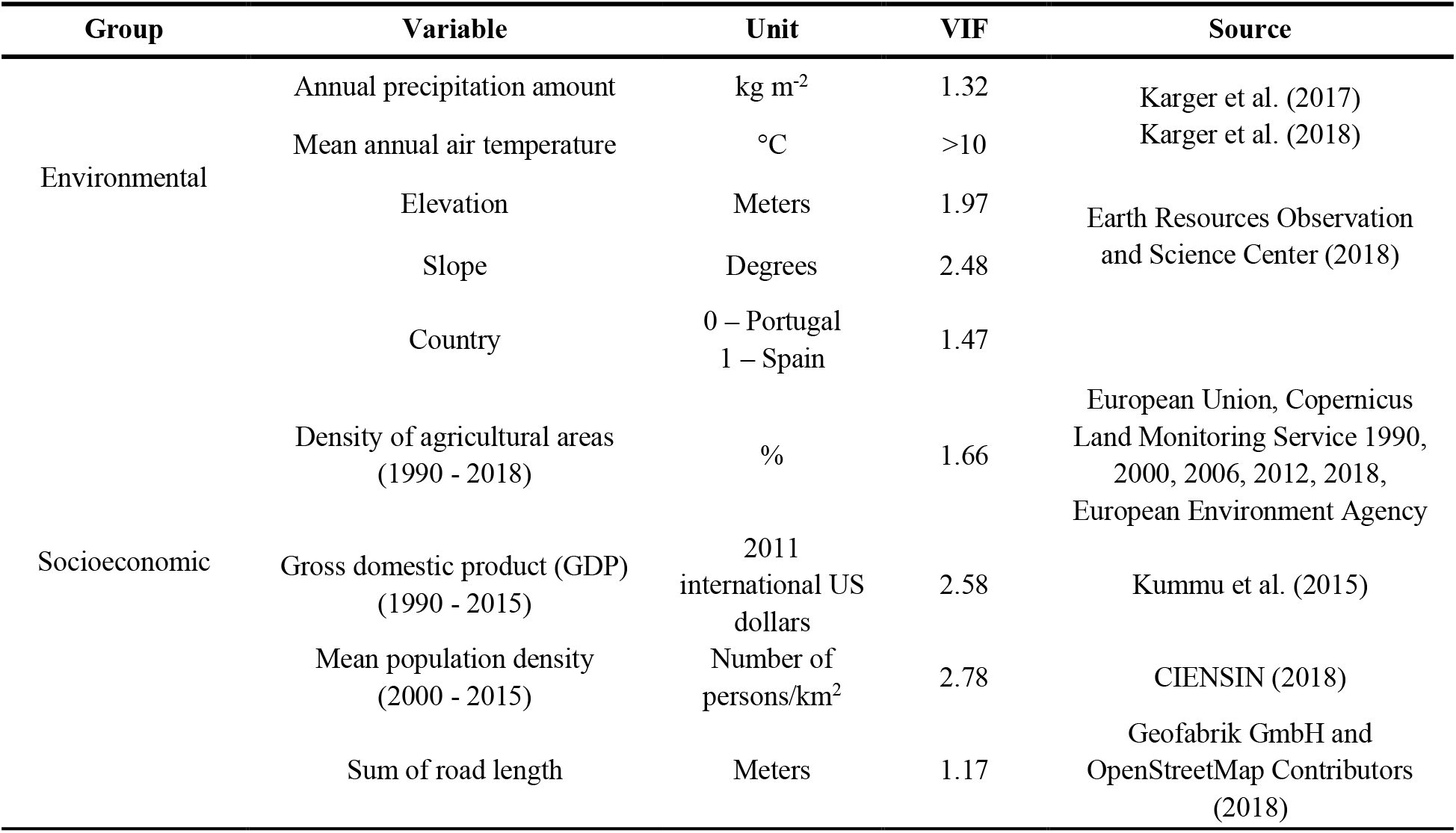
Explanatory environmental and socioeconomic variables tested for detecting spatial differences in LULC dynamics in Iberian peatlands. Only variables with VIF < 10 were included in the multivariate model

Before building the multivariate model, we assessed potential multicollinearity among the variables, using the variation inflation factor (VIF) (Alin 2010). The VIF was calculated the “vifstep” function from the “usdm” R package (Naimi 2022). Only variables with a VIF <10 were considered for modelling (Capinha et al. 2020; Table 2).

#### 2.4.2. Spatial variation explanatory model

We used Boosted Regression Trees (BRT) to assess the relationships between the response variables and the explanatory variables considered (VIF <10, Table 2). BRT are widely used in various research fields, including geography (Cervellini et al. 2021) and ecology (Lemm et al. 2021), and, more particularly, in studies focused on LULC dynamics (Sica et al. 2016). This technique aims to improve the performance of a single classification tree model by sequentially fitting many trees and combining them for prediction (Elith et al. 2008). Four parameters usually need to be defined prior to the calibration of the models (Zhang et al. 2012): the learning rate (LR), tree complexity (TC), the number of trees (NT); and the bag fraction (BF). Following previous recommendations, we used a low value of LR (0.001), an ‘intermediate’ value of TC (5), and a BF between 0.5 and 0.75 (0.66) (Elith et al., 2008). The optimal number of trees was automatically calculated by “gbm.step” function from the “dismo” package (Hijmans et al. 2022) for R.

Because spatial autocorrelation among data points can affect model findings (Crase et al. 2012), we built two BRT models, one not accounting for this issue (the ‘simple’ model); and another where spatial autocorrelation is accounted for by an autocovariate (SAC), representing the spatial variation in residual values from the simple model (Crase et al. 2012). In both models, the response variable was log-transformed (after the addition of a value of one to account for the presence of ‘0’ values) and a Gaussian distribution family was defined for model training (Hijmans et al. 2022). The models were validated using a 5-fold cross-validation procedure, where each fold uses 80% of the data for model training and the remaining 20% for validation. The procedure is repeated for each *s*ubset of the data. The performance of both models was measured using the mean value of the relative absolute error (RAE) of the 5-fold cross-validation sets. RAE values below 100 indicate that the model performs better than a ‘dummy’ predictor, corresponding to the mean of observed values used for evaluation (Capinha et al. 2023).

## 3. Results

### 3.1. Distribution of current peatlands in the Iberian Peninsula

A total of 270 peatland sites were found for Portugal (n=41) and Spain (n=229) (Fig. 1, Supplemental Information Table 1). The Spanish peatlands are mainly located in the North and Central mountain ranges, namely the Toledo Mountains (37%), Central System (30%), and the Iberian System (9%). Considering peatlands located in Portugal, there is a high concentration of these ecosystems in the Northwest mountains (30%) (e.g., the Peneda-Gerês National Park) and in the Serra da Estrela (22%). Nevertheless, peatlands are also found in lowlands in littoral, sublittoral, and rear settings, across the Portuguese Centro region coasts, the Tagus-Sado lower basin, and Southwest Alentejo and Vicentine Coast (15%). Peatlands are under-represented in the south and southeast of Spain, and in the south of Portugal.

### 3.2. LULC change assessment

#### 3.2.1. LULC temporal evolution (1990 - 2018)

Based on the 2000 m radius buffers, from 1990 to 2018, the Iberian peatlands and their surrounding areas were predominantly covered by forest and seminatural areas, with the relative percentage of cover area of this class decreasing negligibly from 84.22%, in 1990, to 83.75%, in 2018 (Supplemental Information Table 2). The second most common class was the agricultural area, also showing a negligible decrease, from 14.70%, in 1990 to 14.59% in 2018. Artificial areas were the third most important class, representing 0.49% of cover in 1990 and 1% in 2018 (i.e., ∼50% of relative increase). Wetlands and water bodies occupy the smallest percentages of area and showed negligible increases (<0.1%). The analysis based on the additional buffer radius (Supplemental Information Table 3 and 4) revealed consistency with the 2000 m buffer results: forest and seminatural areas, and agricultural areas were the classes with the higher relative percentage of cover; the artificial areas registered the greatest relative increase between 1990 and 2018; and the wetlands occupied the least relative percentage of the territory covered by the different buffers.

**Table 3.** Measures (gain, loss, and total change) of LULC transitions, in the 1990 – 2000, 2000 – 2006, 2006 – 2012, and 2012 and 2018 periods

| CLC Classes | Gain | Loss | Total Change |
| --- | --- | --- | --- |
| 1990 - 2000 |  |  |  |
| Artificial Surfaces | 2.48 | 0.27 | 2.75 |
| Agricultural Areas | 7.97 | 9.51 | 17.48 |
| Forest and seminatural areas | 4.51 | 10.00 | 14.51 |
| Wetlands | 0.00 | 0.00 | 0.00 |
| Water bodies | 0.57 | 0.00 | 0.57 |
| 2000 - 2006 |  |  |  |
| Artificial Surfaces | 5.54 | 0.00 | 5.54 |
| Agricultural Areas | 6.80 | 1.18 | 7.98 |
| Forest and seminatural areas | 0.52 | 11.67 | 12.19 |
| Wetlands | 0.00 | 0.00 | 0.00 |
| Water bodies | 0.00 | 0.00 | 0.00 |
| 2006 - 2012 |  |  |  |
| Artificial Surfaces | 6.28 | 0.00 | 6.28 |
| Agricultural Areas | 0.40 | 2.48 | 2.89 |
| Forest and seminatural areas | 0.73 | 4.92 | 5.65 |
| Wetlands | 0.00 | 0.00 | 0.00 |
| Water bodies | 0.00 | 0.00 | 0.00 |
| 2012 - 2018 |  |  |  |
| Artificial Surfaces | 1.46 | 0.85 | 2.31 |
| Agricultural Areas | 4.19 | 1.69 | 5.87 |
| Forest and seminatural areas | 1.69 | 4.79 | 6.48 |
| Wetlands | 0.00 | 0.00 | 0.00 |
| Water bodies | 0.00 | 0.00 | 0.00 |

#### 3.2.2. Main LULC transitions

The transition matrices (Supplemental Information Table 5 A to D) showed that the forest and seminatural areas class has more than 80% of transitions within the same class, in all periods. Concerning transitions between different classes, these occurred mainly from forest and seminatural areas to agricultural areas in 1990 – 2000 (7.97%) and 2012 - 2018 (3.33%) (Supplemental Information Table A and D), and from forest and seminatural areas to artificial areas in 2000 – 2006 (6.80%) and 2006 – 2012 (4.52%) (Supplemental Information Table B and C). The classes that showed the least or no transitions were the wetlands (except for 2000 - 2006, with transitions within the class) and the water bodies (aside from 1990 – 2000, when the class showed gains of 0.57%) (Table 3).

The agricultural areas demonstrated higher cover gains in 1990 to 2000, 2000 to 2006, and 2012 to 2018. In 2006 – 2012, the artificial areas registered the most gains. The area covered by forest and seminatural areas had the highest losses in all periods (Table 3), although, in the first two periods, the loss numbers were higher - 10% in 1990 – 2000 and 11.97% in 2000 – 2006 (Table 3).

#### 3.2.3. Temporal analysis

In all periods, the distributions of the percentage values of the changed area were mainly concentrated between 0% and 10% (Fig. 2). However, compared to other periods, in 1990 – 2000 there were more sites changing 20% to 30% and even 30% to 40% of the total area. In fact, there were significant differences in the distribution of the percentage of changed area between periods (Kruskal-Wallis test, *p* < 0.001), with the period between 1990 and 2000 showing a significantly higher percentage of change than the other periods (Dunn test).

**Fig. 2.**
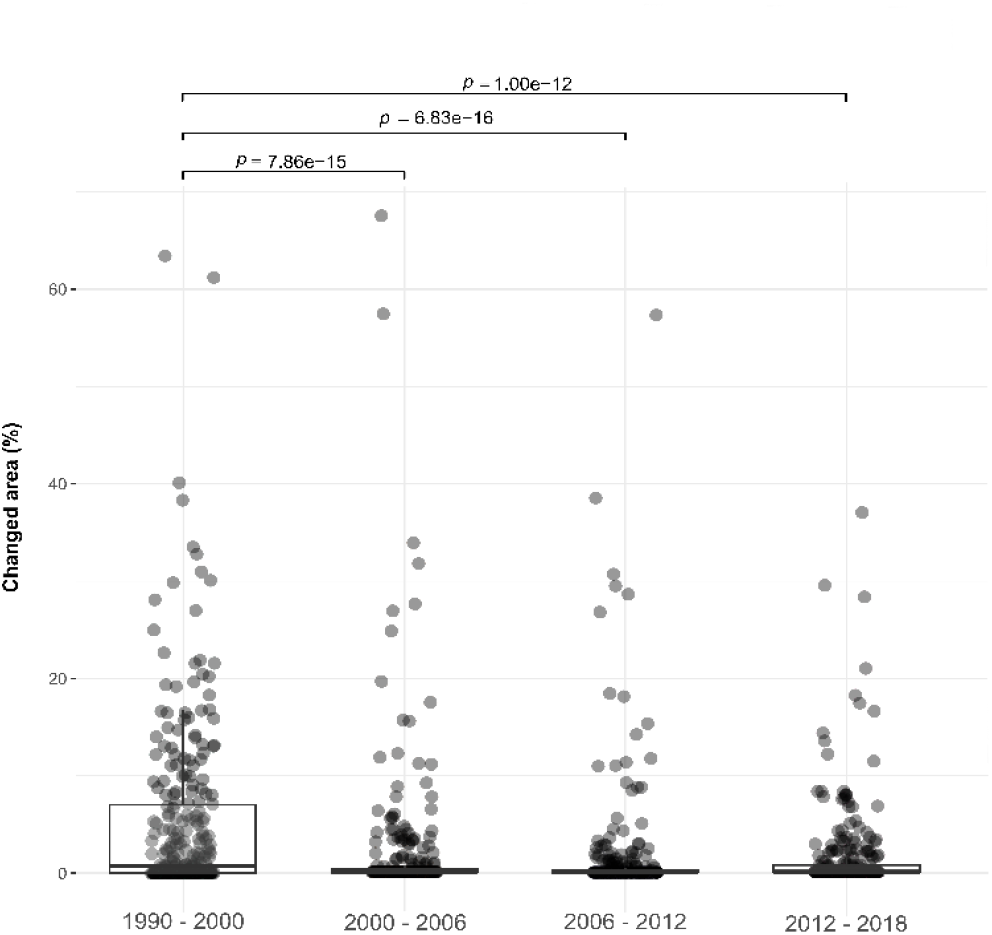
Post-hoc Dunn test results of the differences in the distribution of the percentage of changed area among the studied periods. The *p* < 0.001 between the 1990 – 2000 period and each of the other periods shows that there were significant differences in the percentage of changed area

#### 3.2.4. Area changed between 1990 and 2018

About 93% of the peatland sites experienced an average change in LULC from 0% to 9.51%, with only 7% changing more markedly. The distribution of the percentage of mean changed area between 1990 and 2018 shows an apparent different dynamic in LULC for upland and lowland peatlands (Fig. 3). The strongest LULC changes (9.51% to 38.43% of the total area) were mainly found in ecosystems located in the lowland regions (e.g. fens, coastal lagoons, sublittoral valleys and rear fens and transition mires), mostly in the Portuguese Southwest Alentejo and Vicentine Coast, in the central west coast of Portugal (e.g, Óbidos lagoon), with about 38% of its area changing between 1990 and 2018; and in the Spanish southwest low-altitude area, encircling the Doñana National Park (>20% of alteration). In inland higher regions of Portugal and Spain, we also found several sites where more than 9.51% of the area changed, namely in some sectors of the Iberian mountain chains, namely the northern Iberian System, in the intermediate sector of the Central System, the Toledo Mountains and in the mountains of Northwest Portugal. Notwithstanding, most upland ecosystems showed a mean alteration of less than 9.51%.

**Fig. 3.**
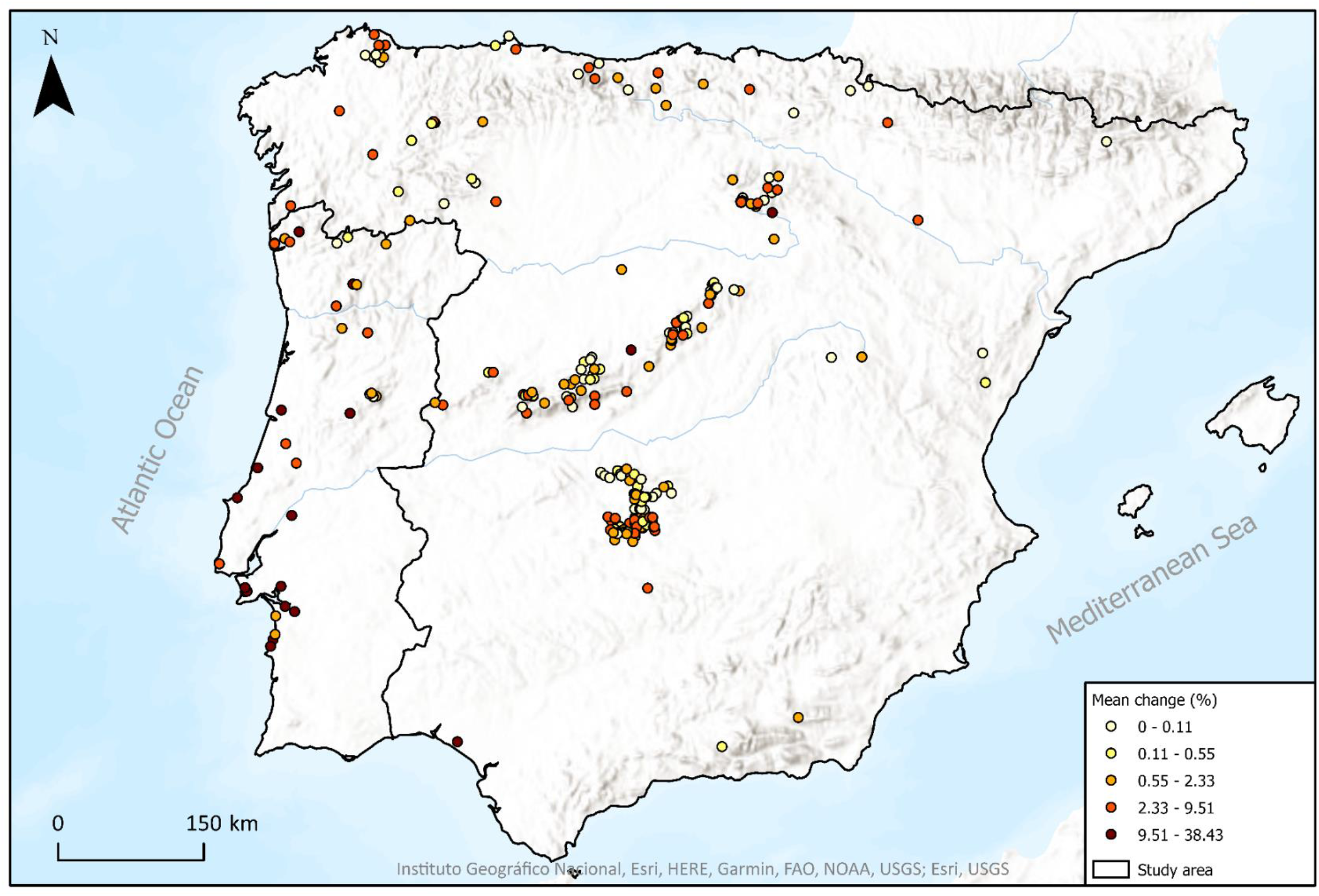
Distribution of the mean changed area, in percentage, between 1990 and 2018, of each peatland sites.

### 3.3. Spatial drivers of LULC dynamics

The BRT model accounting for SAC achieved higher predictive performance than the ‘simple’ model (i.e., lower REA; mean 5-fold = 0.86 ± 0.06), hence we only consider the results of the former. In this model, two predictors showed a relative importance higher than 10% in explaining the spatial distribution of the mean percentage of changed area (Fig. 4): elevation (44.15%) and density of agricultural areas (10.62%).

**Fig. 4.**
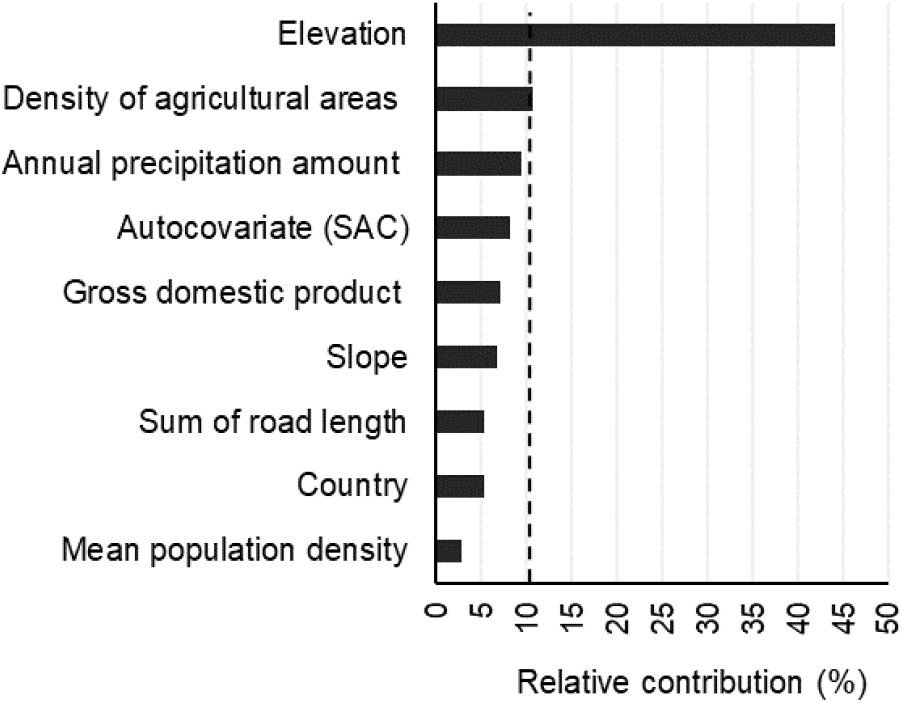
Relative contribution (%) of the predictors of the BRT model. The dashed line marks the threshold at which a predictor contributes to explaining the variation in the dependent variable

Analyzing the partial dependence plot for the elevation predictor (Fig. 5 A), we observed that a higher mean percentage of changed area is associated with sites located at lower altitudes, near the 0 m a.s.l. The probability of change decreases markedly with increasing altitude and stabilizes from around 1300 m a.s.l. Concerning the density of agricultural areas (Fig. 5 B), we observed that locations with farming coverage up to 20% of the total area, thus with less agricultural activity, were associated with a lower mean percentage of change. In peatlands surrounded by 80% or more agricultural areas, the gradient of the response variable decreased substantially.

**Fig. 5.**
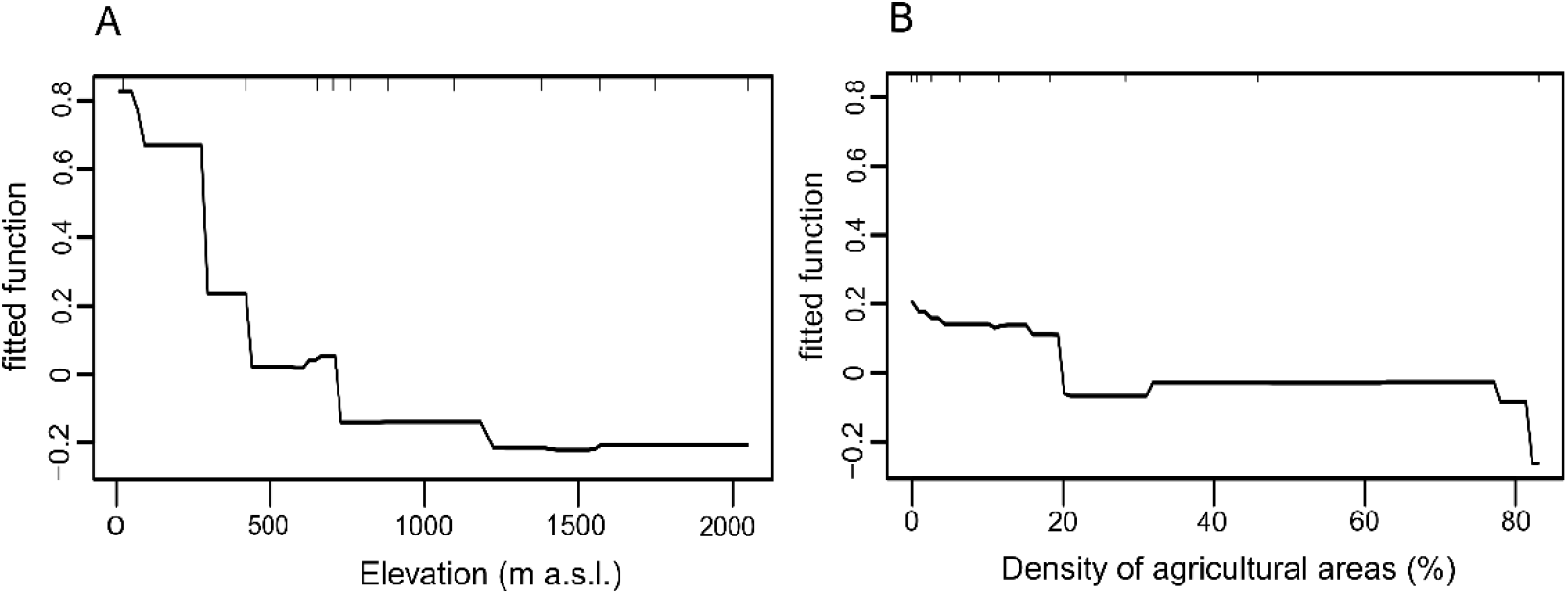
Partial dependence plots for the two most influential variables in the BRT model accounting for spatial autocorrelation. A: Effect of the elevation on the response variable; B: Effect of the density of agricultural areas on the response variable.

## 4. Discussion

Iberian peatlands have undergone major pressures as a result of human-driven LULC changes (Heras Pérez et al. 2017; Mateus et al. 2017; Chico et al. 2019). Our research identified 270 peatlands in the Iberian Peninsula and provided insights into the recent LULC dynamics. Whereas some of these areas showed marked variation in LULC classes over the past ∼30 years, most showed moderate to little change. The propensity for LULC change across sites was mainly driven by altitude variation and agricultural intensity, with higher values for both variables associated with a lower propensity for change.

### 4.1. LULC dynamics in Iberian peatlands

Between 1990 and 2018, the areas of occurrence of peatlands were mainly covered by forest, seminatural, and agricultural areas. For Portugal, these findings are consistent with Meneses et al. (2017), who observed that forest and agricultural areas predominated in the territory between 1995 and 2010. In Spain, according to Corbelle-Rico et al. (2015), the expansion of forests occurred at a higher rate after 1984, which was attributed to the direct influence of the CAP, public afforestation efforts, private initiatives, or natural succession. Forest expansion has occurred countrywide, which can mean that this fact may have also affected peatland areas.

The large extent of forest cover in both countries has been attributed to the decrease in agricultural activities and land abandonment, especially in inland and mountain areas, with impacts on landscape homogenization and species losses (Loidi 2017). Despite the decreasing trend of agricultural areas in the last decades in Portugal (Meneses et al. 2017; Alves et al. 2022), and in areas of steeper slopes in Spain (Fernández-Nogueira & Corbelle-Rico 2020), our results showed that in and around peatlands, gains in the agricultural class were observed between 1990 and 2006, and in 2012-2018. Nevertheless, the main gains of artificial surfaces were consistent with Fernández-Nogueira & Corbelle-Rico (2020) and Alves et al. (2022), showing that the growth of land artificialization mainly affected the coastal areas of the Iberian Peninsula, where many peatlands are located, but also the valley bottoms of mountain areas. The 1990-2000 period stood out for the highest values of percentage of changed area compared to the other periods, suggesting higher LULC dynamics during these years. This dynamic may be attributed to the inclusion of Portugal and Spain in the EU, in 1986, and the subsequent implementation of the CAP (Vidal-Legaz et al. 2013; García-Llamas et al. 2019).

An intriguing result was that the wetlands class covered the least relative percentage of the area, even though our sites included confirmed peatlands. This result may be related to the fact that, since 1970, the surface of natural Mediterranean wetland habitats decreased by 48% (Geijzendorffer et al. 2019). However, problems associated with CLC status layers have been described, some of them directly associated with the wetlands class. For example, limitations such as low resolution and different delineation and classification of wetland polygons can lead to constraints on the use of CLC in wetland monitoring (Ovejero-Campos et al. 2019). Since CHA layers present smaller inconsistencies (García Álvarez & Camacho Olmedo 2023), these layers are recommended to assess LULC changes, rather than CLC status (Büttner et al. 2021).

### 4.2. Degradation status: lowland vs upland

Our BRT model showed that the spatial variable that contributed the most to explaining the gradient of variation in mean change was the elevation. The distribution of changes also indicate that lowland areas have a higher mean changed area than upland areas in high-altitude regions of Portugal and Spain (Fig. 3). These results likely reflect that in the mountain ranges and other remote areas of the Peninsula, rural abandonment and the decrease in human activity (Serra et al. 2008; Lasanta & Vicente-Serrano 2012; Regos et al. 2015) are correlated with substantial increases in forest areas, especially in Spain (Corbelle-Rico et al. 2015), and with the expansion of scrubland or the natural succession processes (Regos et al. 2015). Moreover, it is recognized that mountain peatlands have generally not been reclaimed for agriculture due to the severity of the climate, although they play an important role as summer pastures (Mateus et al. 2017).

Comparatively, flat areas often have more intense human exploitation (Loidi 2017). Nevertheless, our results seem to contradict this expectation, as peatlands located at lower altitudes, which experienced more changes, seem to have a lower density of agricultural areas. However, in lowland peatlands, such as the one near Doñana National Park, previous research has reported the pressure exerted on these ecosystems by intensively-managed farmlands and the sprawl of artificial areas due to economic development pressures (Zorrilla-Miras et al., 2014). Additionally, in some Portuguese lowland peatlands, the reclamation of these areas to rice fields, horticultural plots, and lowland pastures took place in the past few centuries (Mateus et al. 2017). Overall, the observed recent LULC dynamics suggest that upland peatlands are less likely to be degraded by human activity than lowland ecosystems.

### 4.3. Geographical distribution of Iberian peatlands

The current geographical database of peatlands of the Iberian Peninsula is a valuable resource for managers and conservationists and reinforces the geographical patterns identified in previous mapping efforts. We observed that peatlands are more frequent in the Atlantic biogeographical region of both Portugal and Spain, in the Northwest Portuguese Mountains, Galicia, along the Cantabrian range, the Basque Mountains and in Western Navarra, which is consistent with previous research (Martínez-Cortizas et al. 2000; Pontevedra-Pombal et al. 2017). As we move southward, a lower frequency of peatlands is observed and often attributed to the dryer conditions of the Mediterranean biogeographical region (Heras Pérez et al., 2017; Mateus et al., 2017). However, it is possible to observe peat deposits in some coastal wetlands, such as on the Portuguese coast (Mateus et. al. 2017), and in Donãna, southwestern Spain (López-Sáez et al. 2014), which can be related to the favorable peat accumulation conditions, especially in areas that are separated from the sea by sand dunes (Mateus et al. 2017). Also, it is possible to observe peatlands inside the highest valley bottoms and glacial cirques (e.g., Sierra Nevada, south of Spain) (Palma et al. 2017).

Our database shows that peatlands are more widely distributed in the Iberian Peninsula than commonly acknowledged. For example, the Toledo Mountains, in Spain (López-Sáez et al. 2014), and the upland and inland North of Portugal have in fact a higher representation of peatland sites than previously reported. Some of the differences from previous mapping exercises can be attributed to the criteria that were used to develop each database. For example, Heras Pérez et al. (2017) and Mateus et al. (2017) used the criteria of “peatlands areas of international importance”, while Pontevedra-Pombal (2017) mapped Iberian only acid peatlands based on the definition that Loisel et al. (2014) employed to distinguish peat and non-peat material in northern peatlands, which is more adapted to boreal latitudes, where peatlands proliferate. Our integrative approach, which considers both types of sites and is based on the most up-to-date literature and fieldwork, enabled not only a more complete distribution of peatlands on the Iberian Peninsula but also a comprehensive assessment of threats and conservation challenges, leading to a more holistic understanding of these important ecosystems.

## Conclusion

At a time when the role of peatlands in global climate regulation and their importance for biodiversity has become undeniable (Parish et al., 2008; Biancalani & Avagyan, 2014; Tanneberger, Abel, et al., 2021; Tanneberger, Appulo, et al., 2021), the findings of our research represent a key contribution to understanding the extent and conservation status of Iberian peatlands that still withstand degradation factors, namely LULC dynamics. Moreover, our work presents an updated distribution of Iberian peatlands, revealing that many of these ecosystems are located in areas significantly affected by human driven LULC changes in recent decades. While peatlands in high mountain regions generally exhibit lower susceptibility to these changes, we show that peatlands located in lowland areas, especially those in littoral and sublittoral lowlands, have experienced substantial LULC transformations. Therefore, conserving peatlands in lowland regions of the Iberian Peninsula necessitates particular and focused attention within a territory increasingly characterized by significant environmental alterations driven by climate change.

## Contributions

All authors contributed to the study conception and design. Investigation and data collection and analysis were performed by RF and MG. The first draft of the manuscript was written by RF and CC. Supervision was conducted by CC. All authors reviewed and commented on previous versions of the manuscript and read and approved the final manuscript.

