## Supplementary material for "Recent land use and land cover pressures on Iberian peatlands"

**Table 1** Peatland sites in Iberia Peninsula, based on published and unpublished literature and own field work.

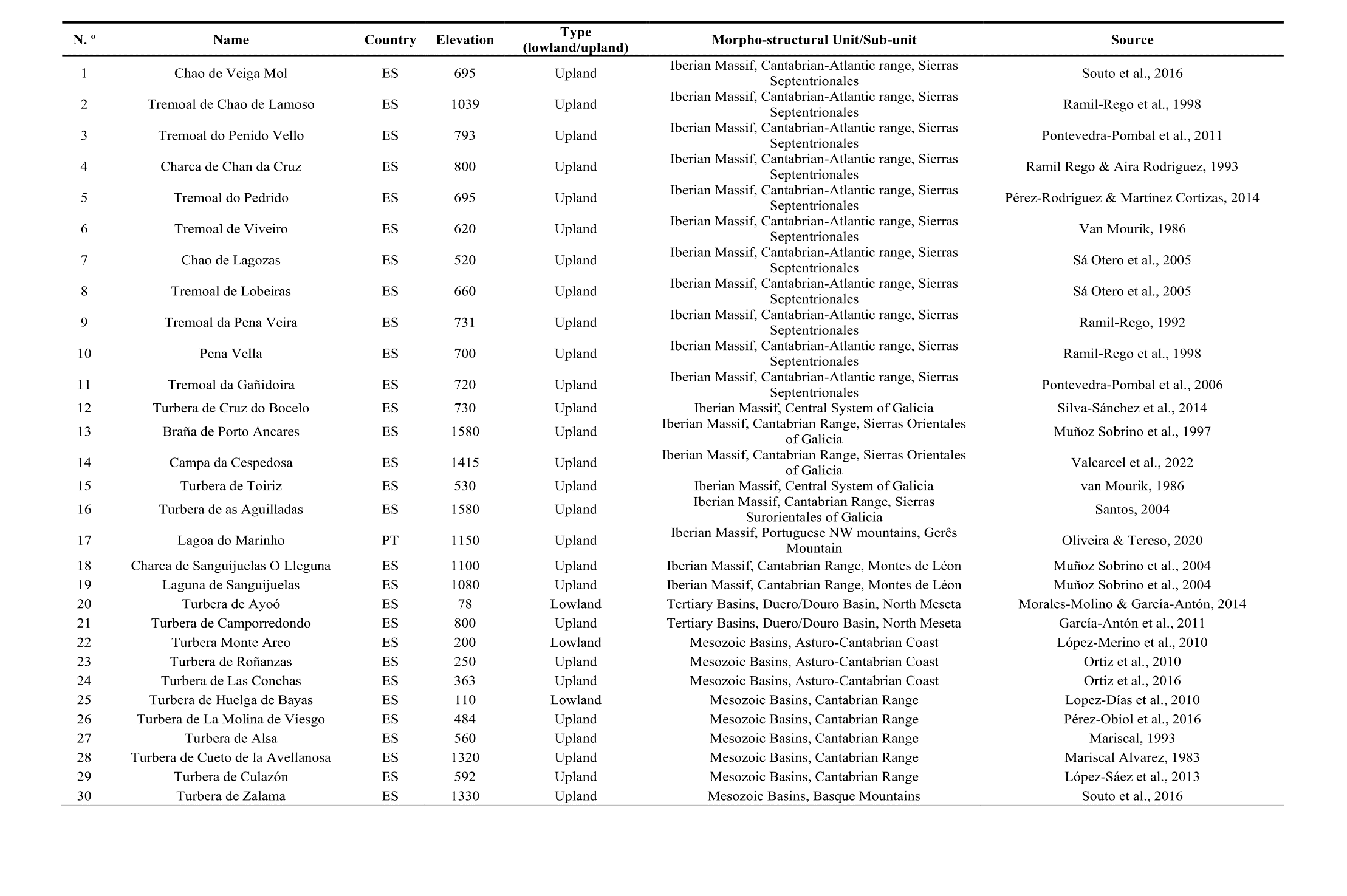

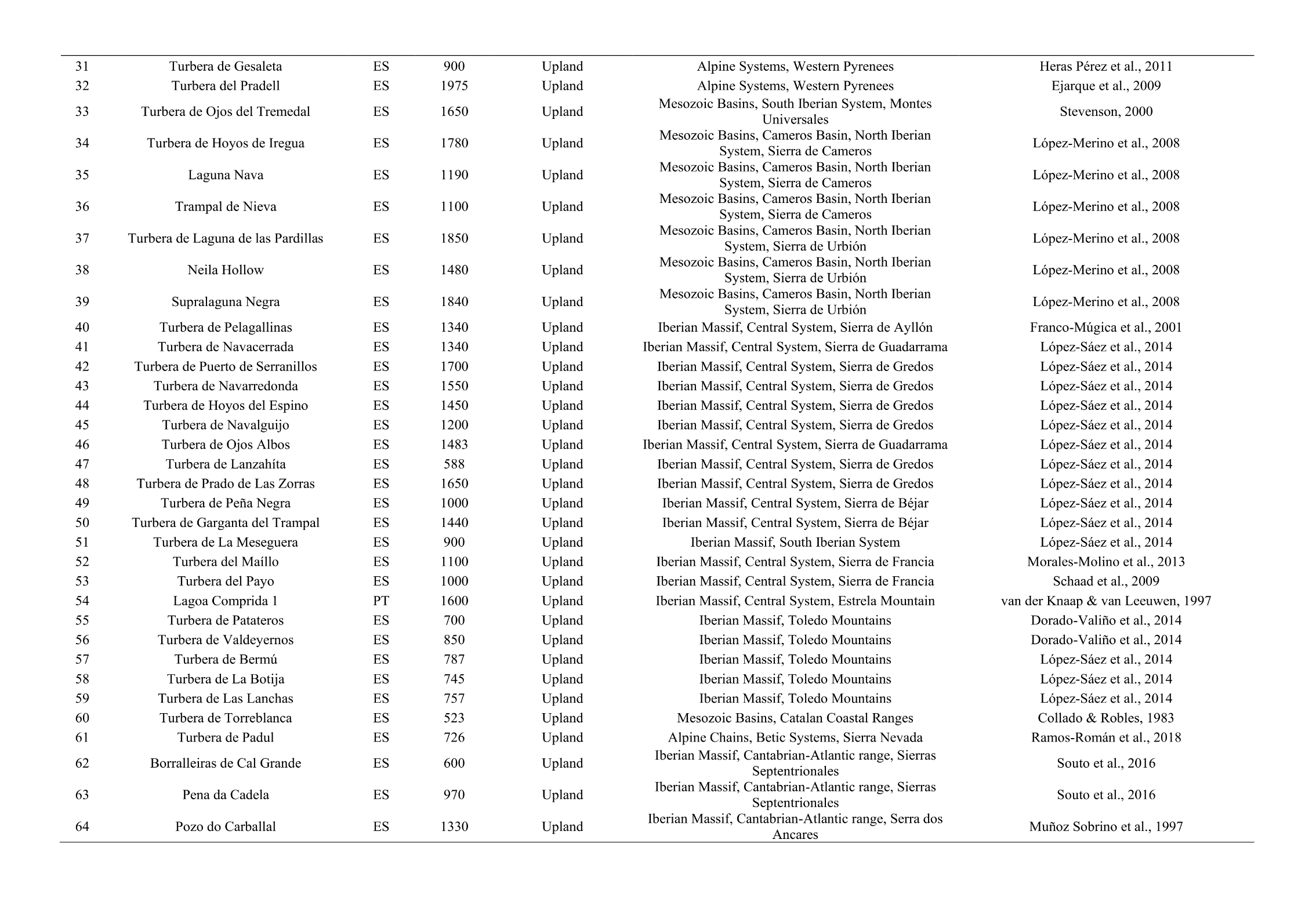

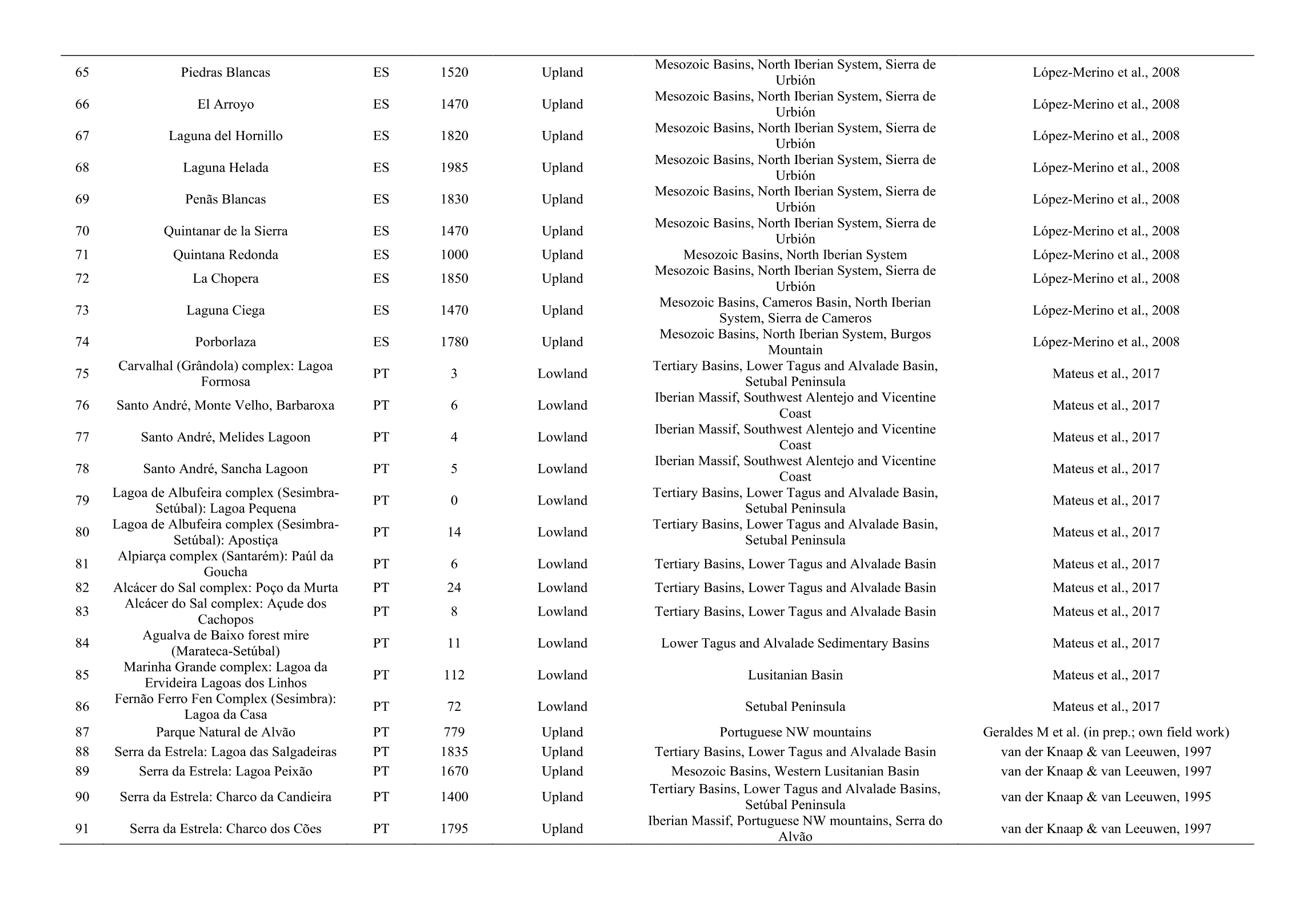

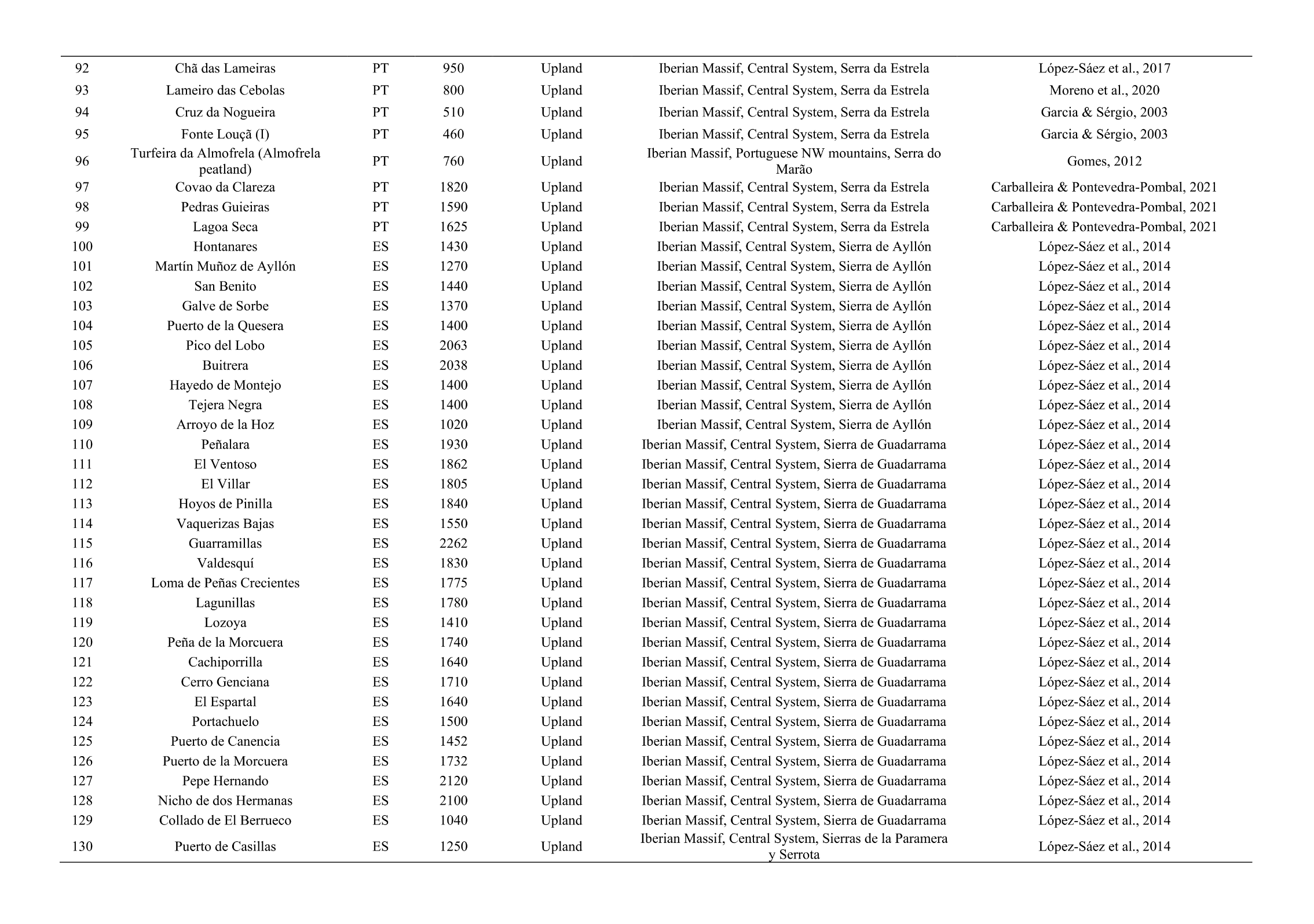

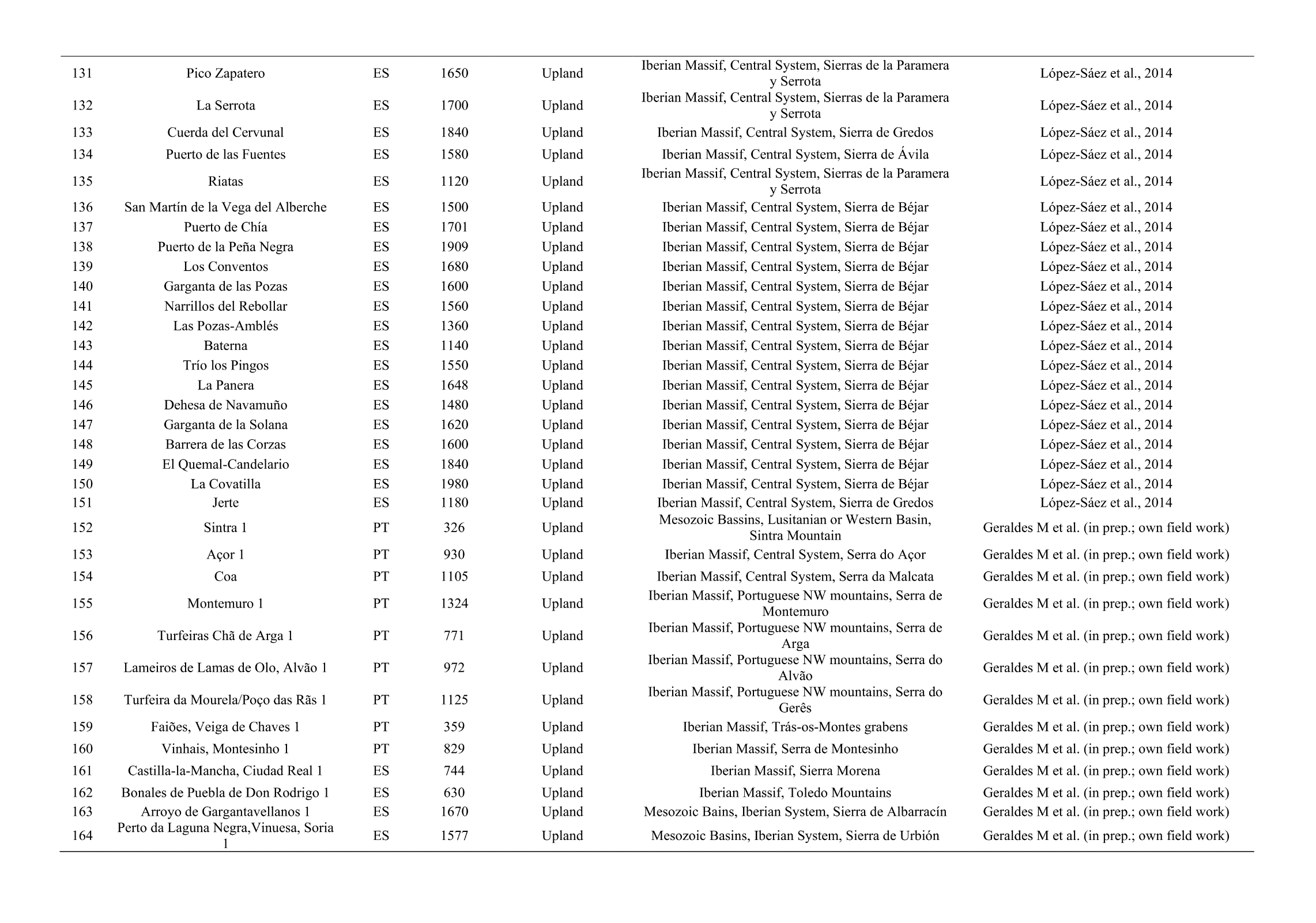

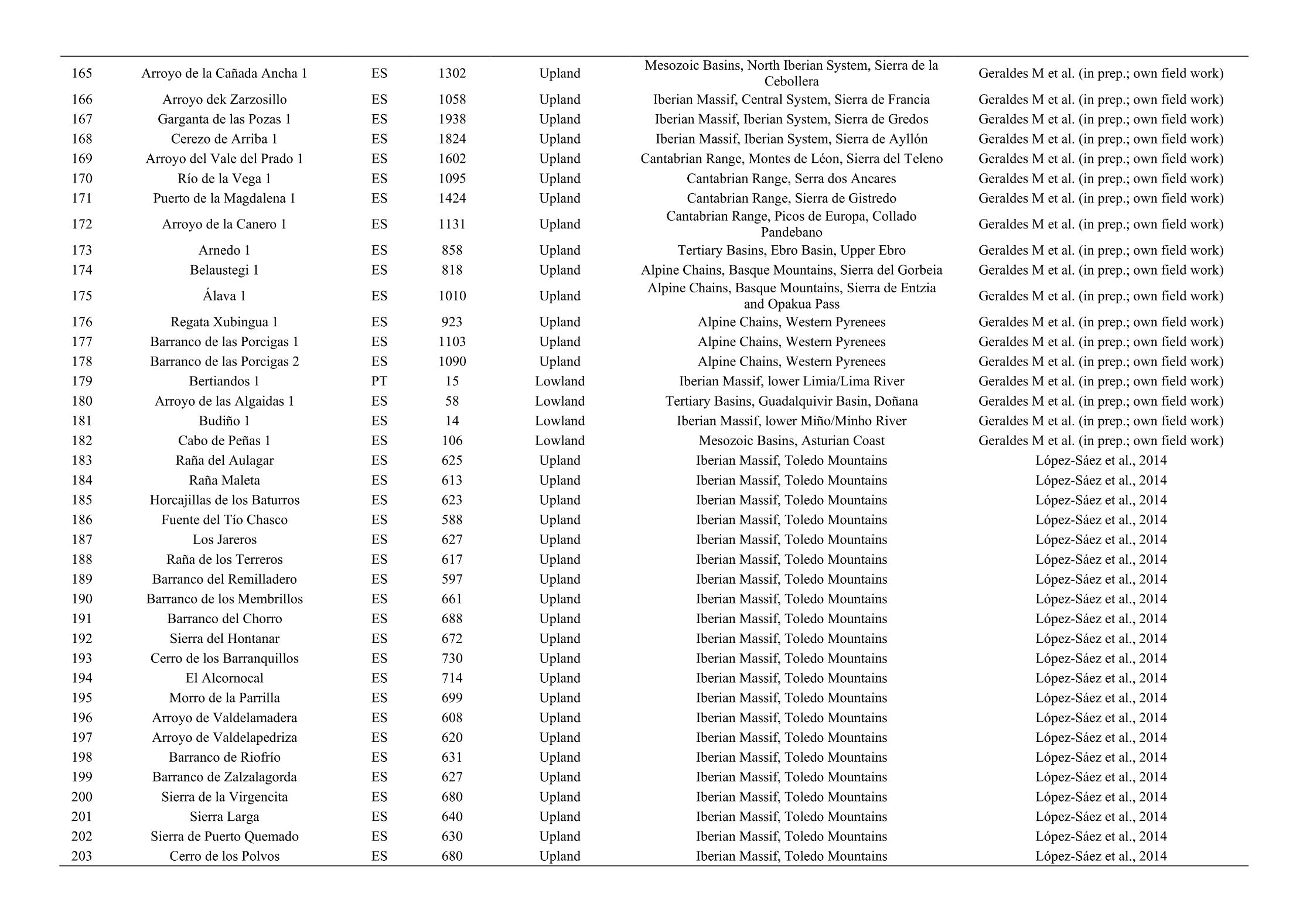

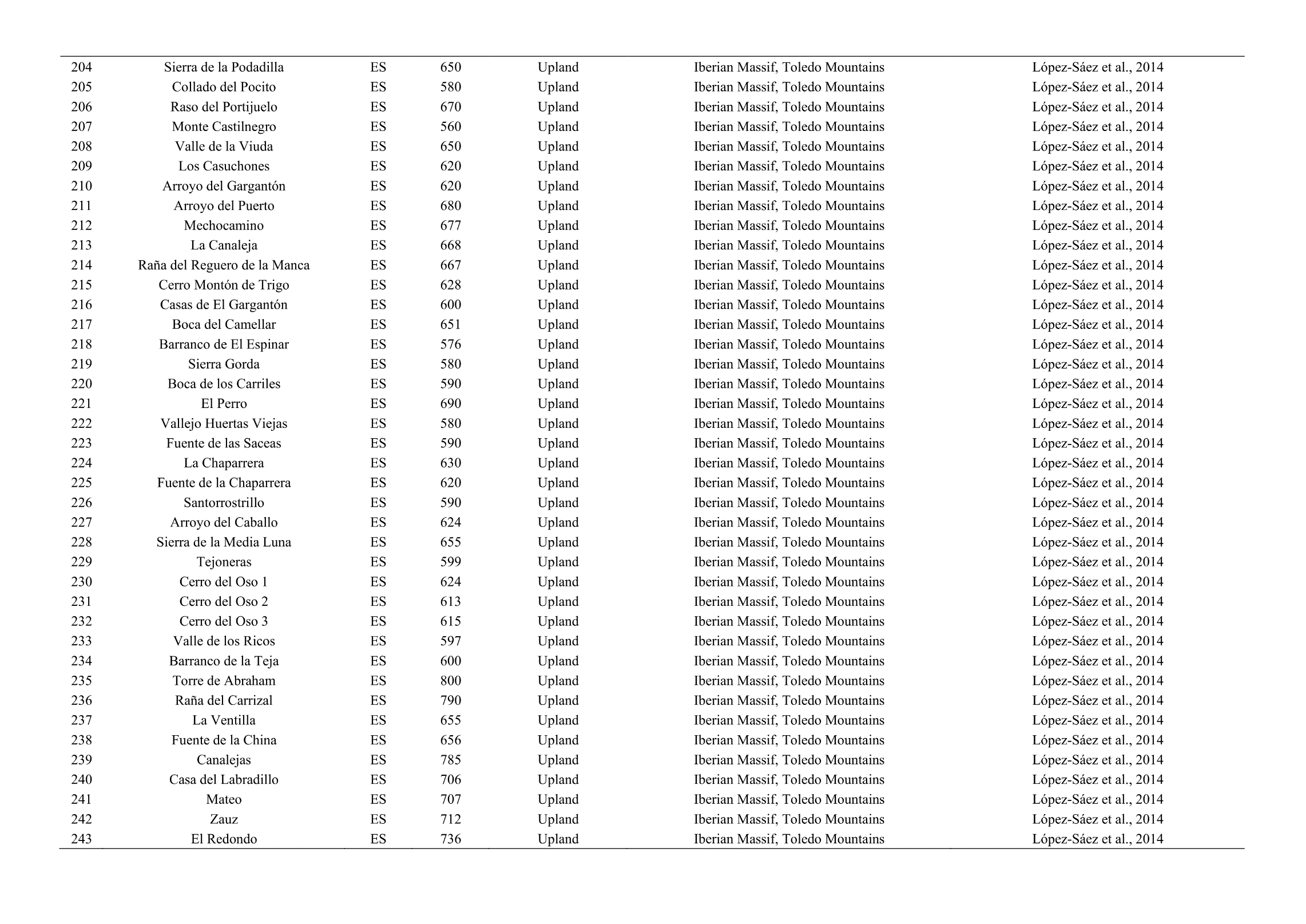

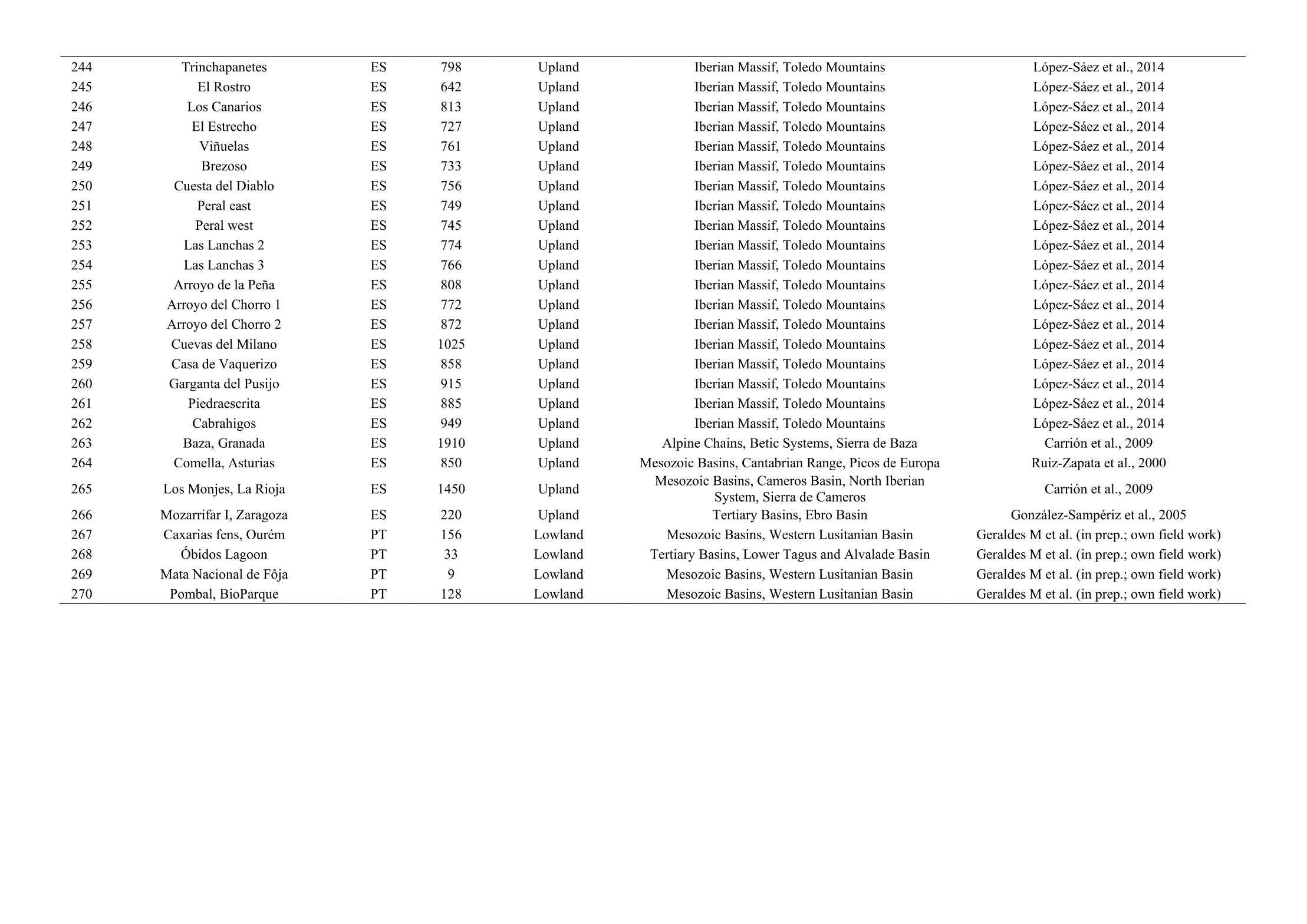

**Table 2** Percentage of relative coverage of the Corine Land Cover level 1 categories in and around peatlands, based on the 2000-m radius buffer.

| **Corine Land Cover**  **(% of coverage, 2000-m radius)** | **1990** | **2000** | **2006** | **2012** | **2018** |
| --- | --- | --- | --- | --- | --- |
| Artificial surfaces | 0.50 | 0.74 | 0.88 | 0.99 | 1.00 |
| Agricultural areas | 14.70 | 14.71 | 14.10 | 14.54 | 14.59 |
| Forest and seminatural areas | 84.22 | 83.92 | 84.37 | 83.81 | 83.75 |
| Wetlands | 0.08 | 0.11 | 0.14 | 0.15 | 0.15 |
| Water bodies | 0.49 | 0.52 | 0.52 | 0.51 | 0.51 |

**Table 3** Percentage of relative coverage of the Corine Land Cover level 1 categories in and around peatlands, based on the 1000-m radius buffer.

| **Corine Land Cover**  **(% of coverage, 1000-m radius)** | **1990** | **2000** | **2006** | **2012** | **2018** |
| --- | --- | --- | --- | --- | --- |
| Artificial surfaces | 0.63 | 0.81 | 0.94 | 0.98 | 0.98 |
| Agricultural areas | 14.37 | 14.37 | 13.66 | 14.29 | 14.44 |
| Forest and seminatural areas | 84.17 | 83.90 | 84.44 | 83.77 | 83.62 |
| Wetlands | 0.21 | 0.27 | 0.26 | 0.27 | 0.27 |
| Water bodies | 0.62 | 0.65 | 0.69 | 0.68 | 0.68 |

**Table 4** Percentage of coverage of each Corine Land Cover level 1 categories in and around peatlands, based on the 4000-m radius buffer.

| **Corine Land Cover**  **(% of coverage, 4000-m radius)** | **1990** | **2000** | **2006** | **2012** | **2018** |
| --- | --- | --- | --- | --- | --- |
| **Artificial surfaces** | 0.64 | 0.96 | 1.15 | 1.24 | 1.26 |
| **Agricultural areas** | 16.40 | 16.24 | 15.48 | 16.03 | 16.08 |
| **Forest and seminatural areas** | 82.31 | 82.10 | 82.66 | 82.04 | 81.97 |
| **Wetlands** | 0.06 | 0.06 | 0.09 | 0.07 | 0.07 |
| **Water bodies** | 0.59 | 0.64 | 0.62 | 0.63 | 0.63 |

**Table 5** Transitions matrices, based on the 2000-m radius buffers. a) From 1990 to 2000; b) From 2000 to 2006; c) From 2006 to 2012; d) From 2012 to 2018.

| **A** | **2000** | | | | | | | |
| --- | --- | --- | --- | --- | --- | --- | --- | --- |
|  | % | Artificial Surfaces | Agricultural Areas | Forest and seminatural areas | Wetlands | Water bodies | Total 1990 | Loss |
| **1990** | Artificial Surfaces | 0 | 0 | 0.05 | 0 | 0.22 | 0.27 | 0.27 |
|  | Agricultural Areas | 0.80 | 4.24 | 4.46 | 0 | 0 | 9.51 | 9.51 |
|  | Forest and seminatural areas | 1.67 | 7.97 | 80.22 | 0 | 0.35 | 90.22 | 10.00 |
|  | Wetlands | 0 | 0 | 0 | 0 | 0 | 0 | 0 |
|  | Water bodies | 0 | 0 | 0 | 0 | 0 | 0 | 0 |
|  | Total 2000 | 2.48 | 12.21 | 84.73 | 0 | 0.57 | 100 |  |
|  | Gain | 2.48 | 7.97 | 4.51 | 0 | 0.57 |  |  |

| **B** | **2006** | | | | | | | |
| --- | --- | --- | --- | --- | --- | --- | --- | --- |
|  | % | Artificial Surfaces | Agricultural Areas | Forest and seminatural areas | Wetlands | Water bodies | Total 1990 | Loss |
| **2000** | Artificial Surfaces | 1.26 | 0 | 0 | 0 | 0 | 1.26 | 0 |
|  | Agricultural Areas | 0.66 | 0.35 | 0.52 | 0 | 0 | 1.53 | 1.18 |
|  | Forest and seminatural areas | 4.87 | 6.80 | 85.53 | 0.00 | 0 | 97.21 | 11.67 |
|  | Wetlands | 0 | 0 | 0 | 0.00 | 0 | 0 | 0 |
|  | Water bodies | 0 | 0 | 0 | 0 | 0 | 0 | 0 |
|  | Total 2000 | 6.80 | 7.15 | 86.05 | 0 | 0 | 100 |  |
|  | Gain | 5.54 | 6.80 | 0.52 | 0 | 0 |  |  |

| **C** | **2012** | | | | | | | |
| --- | --- | --- | --- | --- | --- | --- | --- | --- |
|  | % | Artificial Surfaces | Agricultural Areas | Forest and seminatural areas | Wetlands | Water bodies | Total 1990 | Loss |
| **2006** | Artificial Surfaces | 2.30 | 0 | 0 | 0 | 0 | 2.30 | 0 |
|  | Agricultural Areas | 1.76 | 1.94 | 0.73 | 0 | 0 | 4.42 | 2.48 |
|  | Forest and seminatural areas | 4.52 | 0.40 | 88.24 | 0 | 0 | 93.16 | 4.92 |
|  | Wetlands | 0 | 0 | 0 | 0.12 | 0 | 0.12 | 0 |
|  | Water bodies | 0 | 0 | 0 | 0 | 0 | 0 | 0 |
|  | Total 2000 | 8.58 | 2.34 | 88.96 | 0.12 | 0 | 100 |  |
|  | Gain | 6.28 | 0.40 | 0.73 | 0 | 0 |  |  |

| **D** | | **2018** | | | | | | | |
| --- | --- | --- | --- | --- | --- | --- | --- | --- | --- |
|  |  | % | Artificial Surfaces | Agricultural Areas | Forest and seminatural areas | Wetlands | Water bodies | Total 1990 | Loss |
|  | **2012** | Artificial Surfaces | 2.17 | 0.85 | 0 | 0 | 0 | 3.02 | 0.85 |
|  |  | Agricultural Areas | 0 | 5.89 | 1.69 | 0 | 0 | 7.58 | 1.69 |
|  |  | Forest and seminatural areas | 1.46 | 3.33 | 84.61 | 0 | 0 | 89.40 | 4.79 |
|  |  | Wetlands | 0 | 0 | 0 | 0 | 0 | 0 | 0 |
|  |  | Water bodies | 0 | 0 | 0 | 0 | 0 | 0 | 0 |
|  |  | Total 2000 | 3.62 | 10.08 | 86.29 | 0 | 0 | 100 |  |
|  |  | Gain | 1.46 | 4.19 | 1.69 | 0 | 0 |  |  |

**References Table 1**

Carballeira, R, Pontevedra-Pombal, X (2021) Diversity of testate Amoebae as an indicator of the conservation status of peatlands in Southwest Europe. Diversity 13: 269. <https://doi.org/10.3390/d13060269>

Carrión, J, González-Sampériz, P, López Sáez, J, López García, P, Dupré, M (2009) Quaternary pollen analysis in the Iberian Peninsula: the value of negative results. Internet archaeology 25. <http://intarch.ac.uk/journal/issue25/5/toc.html>

Collado, M, Robles Cuenca, F (1983) Estudio de las asociaciones de moluscos de la Turbera holocena de Torreblanca (Castellón) Mediterránea. Serie de Estudios Geológicos 1: 105-142

Cortizas, M (2014) Preliminary characterization of microbial functional diversity using sole-C-source utilization profiles in Tremoal do Pedrido mire (Galicia, NW Spain). SJSS 4: 158. DOI : 10.3232/SJSS.2014.V4.N2.03

Dorado-Valiño, M, López-Sáez, J, García-Gómez, E, (2014) 21. Patateros, Toledo Mountains (central Spain). Grana 53: 171-173. <https://doi.org/10.1080/00173134.2014.903293>

Ejarque, A, Julià, R, Riera, S, Palet, J, Orengo, H, Miras, Y, Gascón, C (2009) Tracing the history of highland human management in the eastern Pre-Pyrenees: an interdisciplinary palaeoenvironmental study at the Pradell fen, Spain. The Holocene 19: 1241-1255. 10.1177/0959683609345084

Franco-Múgica, F, García-Antón, M, Maldonado, J, Morla, C, Sainz-Ollero, H (2001) Evolución de la vegetación en el sector septentrional del macizo de Ayllón, (Sistema Central). Análisis polínico de la turbera de Pelagallinas (Evolution of vegetation in the northern sector of the Ayllón massif (Central System). Pollen analysis of the Pelagallinas peatland). Anales del Jardín Botánico de Madrid, 59: 113–124.

Garcia, C, Sérgio, C (2003) Novos dados acerca da presença de Bruchia vogesiaca Nestl. ex Schwaegr.(Dicranaceae, Musci) na Serra de Santa Luzia (Minho, Portugal). Portugaliae acta biologica 21: 239-243.

García-Antón, M, Franco-Múgica, F, Morla-Juaristi, C, Maldonado-Ruiz, J (2011) The biogeographical role of Pinus forest on the Northern Spanish Meseta: a new Holocene sequence. Quaternary Science Reviews 30: 757-768.

Gomes, J (2012) The use of digital aerial photography as support for restoration, management and habitat monitoring programmes. Master thesis. Faculty of Sciences. University of Porto. Oporto.

González-Sampériz, P, Valero-Garcés, B L, Carrión, J S, Peña-Monné, J L, García-Ruiz, J, Martí-Bono, C (2005) Glacial and Lateglacial vegetation in northeastern Spain: new data and a review. Quaternary International 140: 4-20. <https://doi.org/10.1016/j.quaint.2005.05.006>

Heras Pérez, P, Infante Sanchez, M, Biurrun Galarraga, I, Campos Prieto, J, Gartziandia Berástegui, A (2011) Tipología, vegetación y estado de conservación de los habitats hidroturbosos del noroeste de Navarra. Acta Botánica Barcinonensia 53: pp 27-45.
https://raco.cat/index.php/ActaBotanica/article/view/252899

López-Días, V, Borrego, A, Blanco, C, Arboleya, M, López-Sáez, J, López-Merino, L (2010) Biomarkers in a peat deposit in Northern Spain (Huelga de Bayas, Asturias) as proxy for climate variation. Journal of Chromatography A, 1217(21), 3538-3546. <https://doi.org/10.1016/j.chroma.2010.03.038>

López-Merino, L, López-Sáez, J, Zapata, M, & García, M (2008) Reconstructing the history of beech (Fagus sylvatica L.) in the north-western Iberian Range (Spain): from Late-Glacial refugia to the Holocene anthropic-induced forests. Review of Palaeobotany and Palynology 152: 58-65. <https://doi.org/10.1016/j.revpalbo.2008.04.003>

López-Sáez, J, Abel-Schaad, D, Alba-Sánchez, F, González-Pellejero, R, Frochoso, M, Allende, F (2013) 20. Culazón, Cantabrian Mountains (northern Spain). Grana 52: 316-318. <https://doi.org/10.1080/00173134.2013.768700>

López-Sáez, J, Abel-Schaad, D, Pérez-Díaz, S, Blanco-González, A, Alba-Sánchez, F, Dorado, M, Ruiz-Zapata, B, Gil-García, M, Gómez-González, C, Franco-Múgica, F (2014) Vegetation history, climate and human impact in the Spanish Central System over the last 9000 years. Quaternary International 353: 98-122. <https://doi.org/10.1016/j.quaint.2013.06.034>

López-Sáez, J A, García-Río, R, Alba-Sánchez, F, García-Gómez, E, Pérez Díaz, S, 2014 Peatlands in the Toledo Mountains (central Spain): characterisation and conservation status. Mires and Peat 15, 1-23. <http://mires-and-peat.net/pages/volumes/map15/map1504.php>.

López-Sáez, J, Figueiral, I, Cruz, D (2017) Palaeoenvironmental and vegetation dynamics in Serra da Nave (Alto Paiva, Beira Alta, Portugal) during the Late Pleistocene and the Holocene. Estudos Pré-Históricos 17, Atas da mesa-redonda “A Pré-história e a Proto-história no Centro de Portugal: avaliação e perspectivas de futuro”. Mangualde, 26 – 27 November 2011, pp. 11-23.

Mariscal Alvarez, B (1983) Estudio polínico de la turbera del Cueto de la Avellanosa, Polaciones (Cantabria). Cadernos do Laboratorio Xeolóxico de Laxe, 1983, 5: 205-226 ISBN: 84-7492-198-8. <http://hdl.handle.net/2183/5858>

Mariscal, B (1993) Variacion de la vegetacion Holocena (4300-280 BP) de Cantabria a traves del analisis polinico de la turbera del Alsa. Estudios Geológicos, 49: 63-68. <https://doi.org/10.3989/egeol.93491-2338>

Mateus, J, Queiroz, P, Joosten, H (2017) Portugal. In Joosten, H, Tanneberger, F, Moen, A (ed) Mires and peatlands of Europe. Mires and peatlands of Europe, Schweizerbart, Stuttgart, pp. 572–579.

Morales-Molino, C, García-Antón, M, 2014 Vegetation and fire history since the last glacial maximum in an inland area of the western Mediterranean Basin (Northern Iberian Plateau, NW Spain). Quaternary Research, 81(1), 63-77. <https://doi.org/10.1016/j.yqres.2013.10.010>

Morales-Molino, C, García-Antón, M, Postigo-Mijarra, J, Morla, C (2013) Holocene vegetation, fire and climate interactions on the westernmost fringe of the Mediterranean Basin. Quaternary Science Reviews, 59, 5-17. <https://doi.org/10.1016/j.quascirev.2012.10.027>

Moreno, F, Moreno, J, Fatela, F, Guise, L, Vieira, C, Leira, M (2020) Bromine biogeodynamics in the NE Atlantic: A perspective from natural wetlands of western Portugal. Science of The Total Environment 722: 137649. <https://doi.org/10.1016/j.scitotenv.2020.137649>

Mugica, F, Antón, M, Ruiz, J, Juaristi, C, Ollero, H (2001) Evolución de la vegetación en el sector septentrional del macizo de ayllón (sistema central). Análisis polínico de la turbera de pelagallinas. Anales del Jardín Botánico de Madrid 59: 113-124. Consejo Superior de Investigaciones Científicas.

Muñoz Sobrino, C, Ramil-Rego, P, & Rodríguez Guitián, M (1997) Upland vegetation in the north-west Iberian peninsula after the last glaciation: forest history and deforestation dynamics. Vegetation History and Archaeobotany 6: 215-233. <https://doi.org/10.1007/BF01370443>

Muñoz Sobrino, C, Ramil-Rego, P, Gómez-Orellana, L (2004) Vegetation of the Lago de Sanabria area (NW Iberia) since the end of the Pleistocene: a palaeoecological reconstruction on the basis of two new pollen sequences. Vegetation History and Archaeobotany 13: 1–22. <https://doi.org/10.1007/s00334-003-0028-1>

Oliveira, C, Tereso, J (2020) Dinâmicas de vegetação no final do Pleistocénico e início do Holocénico no atual território português. Arqueologia & História 70: 133-146. <http://hdl.handle.net/10451/44678>

Ortiz, J, Borrego, Á, Gallego, J, Sánchez-Palencia, Y, Urbanczyk, J, Torres, T, Domingo, L, Estébanez, B (2016) Biomarkers and inorganic proxies in the paleoenvironmental reconstruction of mires: the importance of landscape in Las Conchas (Asturias, Northern Spain). Organic geochemistry, 95, 41-54. <https://doi.org/10.1016/j.orggeochem.2016.02.009>

Ortiz, J, Gallego, J, Torres, T, Díaz-Bautista, A, Sierra, C (2010) Palaeoenvironmental reconstruction of Northern Spain during the last 8000 cal yr BP based on the biomarker content of the Roñanzas peat bog (Asturias).  Organic Geochemistry 41: 454-466. <https://doi.org/10.1016/j.orggeochem.2010.02.003>

Pérez-Obiol, R, García-Codron, J C, Pelachs, A, Pérez-Haase, A, Soriano, J (2016) Landscape dynamics and fire activity since 6740 cal yr BP in the Cantabrian region (La Molina peat bog, Puente Viesgo, Spain). Quaternary Science Reviews 135: 65-78.

Pérez-Rodríguez, M, Cortizas, M (2014) Preliminary characterization of microbial functional diversity using sole-C-source utilization profiles in Tremoal do Pedrido mire (Galicia, NW Spain). SJSS 4: 158. <https://doi.org/10.1016/j.quascirev.2016.01.021>

Pontevedra-Pombal, X, Muñoz, J, García-Rodeja, E, Cortizas, A (2006) Mountain mires from Galicia (NW Spain). Developments in Earth Surface Processes 9: 85-109. <https://doi.org/10.1016/S0928-2025(06)09004-3>

Pontevedra-Pombal, X, Mighall, T, Nóvoa-Muñoz, J, Peiteado-Varela, E, Rodríguez-Racedo, J, García-Rodeja, E, Martínez-Cortizas, A (2013) Five thousand years of atmospheric Ni, Zn, As, and Cd deposition recorded in bogs from NW Iberia: prehistoric and historic anthropogenic contributions. Journal of Archaeological Science 40: 764-777. <https://doi.org/10.1016/j.jas.2012.07.010>

Ramil-Rego, P (1992) La vegetación cuaternaria de las sierras septentrionales de Lugo a través del análisis polínico (Quaternary vegetation of the northern mountains of Lugo according to pollen analysis). PhD thesis, University of Santiago de Compostela, 356 pp.

Ramil Rego, P, Aira Rodriguez, M (1993) Análisis polínico de la turbera de la Charca do Chan da Cruz (Ferreira do Valadouro, Lugo. NO de España). Ecologia mediterranea 19: 71-78. <https://doi.org/10.3406/ecmed.1993.1723>

Ramil-Rego, P, Muñoz-Sobrino, C, Rodríguez-Guitián, M, Gómez-Orellana, L (1998) Differences in the vegetation of the North Iberian Peninsula during the last 16,000 years. Plant Ecology 138: 41-62. <https://doi.org/10.1023/A:1009736432739>

Ramos-Román, M, Jiménez-Moreno, G, Camuera, J, García-Alix, A, Anderson, R, Jiménez-Espejo, F, Sachse, D, Toney, J, Carrión, J, Webster, C, Yanes, Y (2018) Millennial-scale cyclical environment and climate variability during the Holocene in the western Mediterranean region deduced from a new multi-proxy analysis from the Padul record (Sierra Nevada, Spain). Global and Planetary Change 168: 35-53. <https://doi.org/10.1016/j.gloplacha.2018.06.003>

Sá Otero, M, Porto, A, Losada, E (2005) A study of the post-glacial vegetation in" Montes do Buio"(NW Spain). Lagascalia 25: 91-114.

Santos, L (2004) Late Holocene forest history and deforestation dynamics in the Queixa Sierra, Galicia, northwestern Iberian Peninsula. Mountain Research and Development, 24(3), 251-257. [https://doi.org/10.1659/0276-4741(2004)024[0251:LHFHAD]2.0.CO;2](https://doi.org/10.1659/0276-4741(2004)024%5b0251:LHFHAD%5d2.0.CO;2)

Schaad, D, Hernández, A, López Sáez, J, Pulido Díaz, F, López Merino, L, Martínez Cortizas, A (2009) Evolución de la vegetación en la Sierra de Gata (Cáceres-Salamanca, España) durante el Holoceno reciente. Implicaciones biogeográficas. <http://hdl.handle.net/10261/93749>

Silva-Sánchez, N, Martínez Cortizas, A, & López-Merino, L (2014) Linking forest cover, soil erosion and mire hydrology to late-Holocene human activity and climate in NW Spain. The Holocene 24: 714-725. DOI: 10.1177/0959683614526934

Souto, M, Castro, D, Pontevedra-Pombal, X, Garcia-Rodeja, E, Fraga, M (2016) Characterisation of Holocene plant macrofossils from North Spanish ombrotrophic mires: vascular plants. Mires and Peat 18: 1-21. DOI: 10.19189/MaP.2016.OMB.236

Stevenson, A (2000) The Holocene forest history of the Montes Universales, Teruel, Spain. The Holocene, 10: 603-610.

Valcarcel, M, Vázquez-Rodríguez, A, Pontevedra-Pombal, X (2022) Inestabilidad de ladera natural e inducida asociada a grandes movimientos en masa durante el Pleistoceno-Holoceno en la Serra dos Ancares (NW de la Península Ibérica). Anales de Geografía de la Universidad Complutense 42: 31. <https://dx.doi.org/10.5209/aguc.81806>

Van der Knaap, W, Van Leeuwen, J (1995) Holocene vegetation succession and degradation as responses to climatic change and human activity in the Serra de Estrela, Portugal. Review of Palaeobotany and Palynology 89: 153-211. <https://doi.org/10.1016/0034-6667(95)00048-0>

Van der Knaap, W, Van Leeuwen, J (1997) Late Glacial and early Holocene vegetation succession, altitudinal vegetation zonation, and climatic change in the Serra da Estrela, Portugal. Review of Palaeobotany and Palynology 97: 239-285. <https://doi.org/10.1016/S0034-6667(97)00008-0>

Van Mourik, J (1986) Pollen Profiles of Slope Deposits in the Galcian Area (NW Spain). Netherlands Geographical Studies 12, Koninklijk Nederlands Aardrijkskundig Genootschap / Fysisch Geografisch en Bodemkundig Laboratorium van de Universiteit van Amsterdam, Amsterdam, 171 pp.
